# SalmonAct Deciphers Transcription Factor Regulatory Activity in *Salmonella* Transcriptomics

**DOI:** 10.1101/2025.08.22.671797

**Authors:** Marton Olbei, Balazs Bohar, Robert A. Kingsley, Tamas Korcsmaros

**Affiliations:** Division of Digestive Diseases, Department of Metabolism, Digestion and Reproduction, Imperial College London, United Kingdom; Earlham Institute, Norwich, United Kingdom; Quadram Institute, Norwich, United Kingdom

## Abstract

Foodborne *Salmonella enterica* infection remains a major public health threat due to its prevalence in food, ease of transmission, and increasing antibiotic resistance. Consequently, understanding the molecular mechanisms underlying *Salmonella* pathogenesis is crucial for guiding novel diagnostic, preventative, and therapeutic approaches. High-throughput transcriptomic technologies are now often employed in *Salmonella* research to quantify how gene expression changes in response to various conditions or mutations. But due to the high dimensionality of this data, resulting from the comparison of the expression of thousands of genes from multiple strains or culture conditions with complex interactions, interpretation remains a demanding task. To address this issue, we present SalmonAct, a comprehensive signed and directed prior knowledge resource for inferring transcription factor activities in *Salmonella*. This new resource expands the toolkit available for *Salmonella* functional analysis. Built as an extension of the SalmoNet2 database, SalmonAct can be used to infer the activity of 191 transcription factors in 5991 regulatory interactions based on publicly available interaction and regulatory knockout data. SalmonAct enhances the interpretation of highly-dimensional transcriptomic data by identifying both highly influential and minimally active transcription factors that drive the observed expression state. SalmonAct aids in bridging the gap between model and non-model organisms’ functional analysis, and together with the SalmoNet2 resource, can be used for further downstream data analyses.

**Importance:** While modern high-throughput transcriptomics technologies provide effective ways of studying *Salmonella enterica*, the complexity of these datasets makes interpretation challenging. SalmonAct provides a new computational resource to connect gene expression changes with the activity of transcription factors, key regulators of bacterial adaptation and virulence. SalmonAct can help researchers progress from the analysis of gene expression profiles towards a clearer understanding of the regulatory programs driving the behaviour of the pathogen. By integrating prior knowledge of transcription factor - target gene relationships, SalmonAct contributes to uncovering novel mechanisms of bacterial pathogenesis, and supports the development of new approaches to combat *Salmonella* infections.

## Introduction

*Salmonella enterica*, a globally significant pathogen, causes a diverse range of illnesses, from mild gastroenteritis to life-threatening systemic infections (*1*). It poses a significant public health challenge due to its widespread presence in the food supply, the ease with which it can spread, and the increasing prevalence of antibiotic-resistant strains (*2, 3*). Therefore, a comprehensive understanding of the molecular mechanisms underlying *Salmonella* pathogenesis is crucial for developing novel diagnostic, preventive, and therapeutic approaches (*4*). High-throughput transcriptomic technologies play a pivotal role in identifying these mechanisms, by revealing how gene expression changes in response to environments encountered by the pathogen during its lifecycle, enabling targeted studies of key regulators and pathways (*5*).

As the cost of data generation continues to decrease, the volume of publicly available transcriptomic datasets is rapidly growing, making it increasingly important to develop methods for extracting functional information from these results (*6, 7*). While analysing RNA level changes across conditions can provide valuable insights into a system’s state, the high complexity of these data can make interpretation challenging. Leveraging prior knowledge networks (PKNs), which represent known, experimentally validated relationships within a system, reduces data complexity and enhances statistical power. By aggregating signals from known target genes, these methods estimate transcription factor influence more robustly than by examining the expression of singular genes (*8*). Although prior knowledge resources are increasingly available for well-characterised model organisms including *E. coli*, they remain limited for other important pathogens such as *Salmonella enterica (4)*.

One such application of PKNs is the inference of transcription factor (TF) activity. TF activity is the influence a TF exerts on the transcriptome, evaluated through the expression of the TF’s target genes. TFs can bind specific DNA sequences to either activate or repress transcription. Their overall activity, or influence, can be inferred by examining whether their known target genes are up- or downregulated, in line with the TF’s mode of regulation (inhibition or stimulation). By assessing whether sets of target genes are up- or downregulated in a given dataset, we gain a more complete view of TF function, rather than relying solely on the TF’s own differential expression status. In other words, this approach aggregates signals from multiple genes to identify whether a TF’s regulatory influence is broadly active or inactive under the condition of interest, based on the behaviour of the TF’s known target genes.

PKNs for non-model organisms such as *Salmonella* are rare, and tend to focus on different molecular mechanisms. For example, the KEGG (*9*) and BioCyc (*10*) databases collate *Salmonella*-specific metabolic pathways, the STRING database (*11*) contains protein associations, but neither of them contain gene regulatory interactions. Specifically for gene regulatory networks, the CollecTF (*12*) and PRODORIC (*13*) databases contain signed, directed, experimentally validated *Salmonella* interactions, however their coverage is limited. Other resources, such as RegPrecise (*14*) contain a more expansive, manually curated set of regulatory interactions, and databases such as SalComRegulon (*15*) collate the effect of regulatory knockouts on the *Salmonella* transcriptome. Previously we have developed SalmoNet2 (*16*), a multi-layered *Salmonella* interaction resource, that simultaneously integrates protein-protein, transcriptional regulatory, and metabolic interactions to provide an enhanced view of *Salmonella* biology. Although SalmoNet2 includes a comprehensive gene regulatory layer for 20 *Salmonella* strains, these gene regulatory interactions are not signed, and thus cannot be used for transcription factor activity inference.

Here, we introduce SalmonAct, a signed and directed prior knowledge resource specifically developed for *Salmonella* transcription factor activity inference. SalmonAct is designed to work alongside established methods for inferring transcription factor influences. By combining transcription factor activities with transcriptomics data, this integration enables researchers to gain a deeper understanding of the mechanisms controlling *Salmonella* biology.

## Results

### A novel prior knowledge network for *Salmonella*

To collate the signed and directed interactions between transcription factors and their regulated target genes, we integrated a prior knowledge network (PKN) from multiple information sources. Data were imported from the CollecTF, PRODORIC, RegPrecise and RegulonDB databases, and regulatory perturbation data from the SalComRegulon dataset (*12*–*15, 17*). The resulting PKN contains 5991 signed and directed regulatory interactions for 191 transcription factors. 44% of interactions are inhibitory, and 56% are stimulatory. For further details on PKN construction please see the Methods section.

The PKN can be accessed directly from the SalmoNet2 website (http://salmonet.org/), and is provided in a format that is directly accessible by applicable methods, such as decoupleR. DecoupleR is an R package and python library developed to extract biological activities from omics data, with a multitude of methods, including their consensus decisions (*8*).

### Inferring transcription factor activities using the SalmonAct network

To explore the potential of SalmonAct, we analysed two *Salmonella enterica* transcriptomics datasets and determined the least and most active transcription factors in each case. In the first use case, we analysed data for genes regulated by the OmpR regulon identified by culturing wild-type and *ompR* mutant *S*. Typhimurium strain SL1344 to mid-exponential phase (*18*). We determined the differentially expressed genes between the two groups (SL1344 ompR /SL1344 wildtype), and analysed the results with SalmonAct and decoupleR. OmpR was found to be the least active transcription factor, as could be expected, following the knockout experiment (Figure 1A). The master regulator *crp* was found to be the most active transcription factor. Crp fills a central role in the physiology of *Salmonella*, integrating environmental signals through intracellular cAMP levels, which it binds as a cofactor (*19*). Figure 1B shows the expression levels of the target genes of OmpR following the knockout, illustrating how they contribute to its low activity score. Since targets OmpR normally activates are downregulated, and many of the targets it normally suppresses are upregulated, in the absence of OmpR, the TF achieves a low score, and is therefore considered inactive.

**Figure 1:**
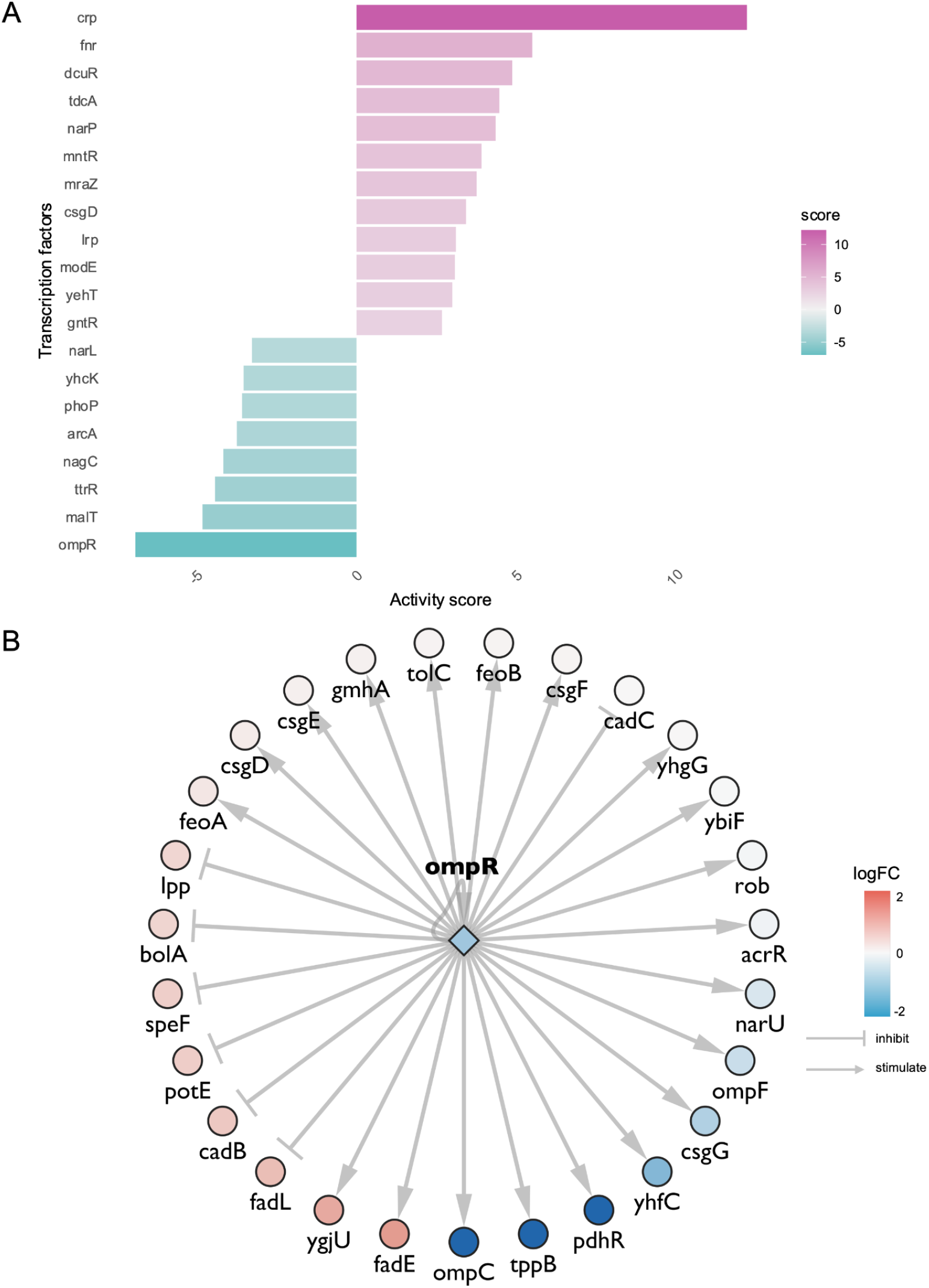
Transcription factor activity analysis on an ompR knockout dataset. A: Bar charts depicting the most and least active transcription factors in the ompR knockout vs wild-type comparison. B: OmpR targets and their differential expression levels, illustrating how the mode of regulation aligns with the observed behaviour of its target genes. Because many genes normally activated by OmpR are downregulated and many that it represses are upregulated, its inferred activity is low.

In the second set of analysed transcriptomics data, we analysed the differential expression of genes during log phase and stationary phase, in a culture of *S*. Typhimurium strain 14028S (*20*). Figure 2A shows the ranking of transcription factor activities. The highest scoring transcription factor, CsgD, is the master regulator of biofilm development in *Salmonella*. The high activity is to be expected, as cells transition from motile cells to the stationary phase, and begin biofilm formation (*21*). YncC (also known as McbR), the third-most active transcription factor, similarly stimulates biofilm formation in *E. coli* (*22*). The least active transcription factor, fis, fills a central role in regulating metabolic and type III secretion factors, targeting many of the SPI-1 and SPI-2 genes (*23, 24*). Figure 2B shows the differential expression results on a volcano plot, with YncC, CsgD and fis highlighted. Neither transcription factors are differentially expressed, but by involving data from the collective behaviour of their target genes, we can get a clearer idea of how the system as a whole is behaving instead of having to rely on the differential expression status of individual transcription factors.

**Figure 2:**
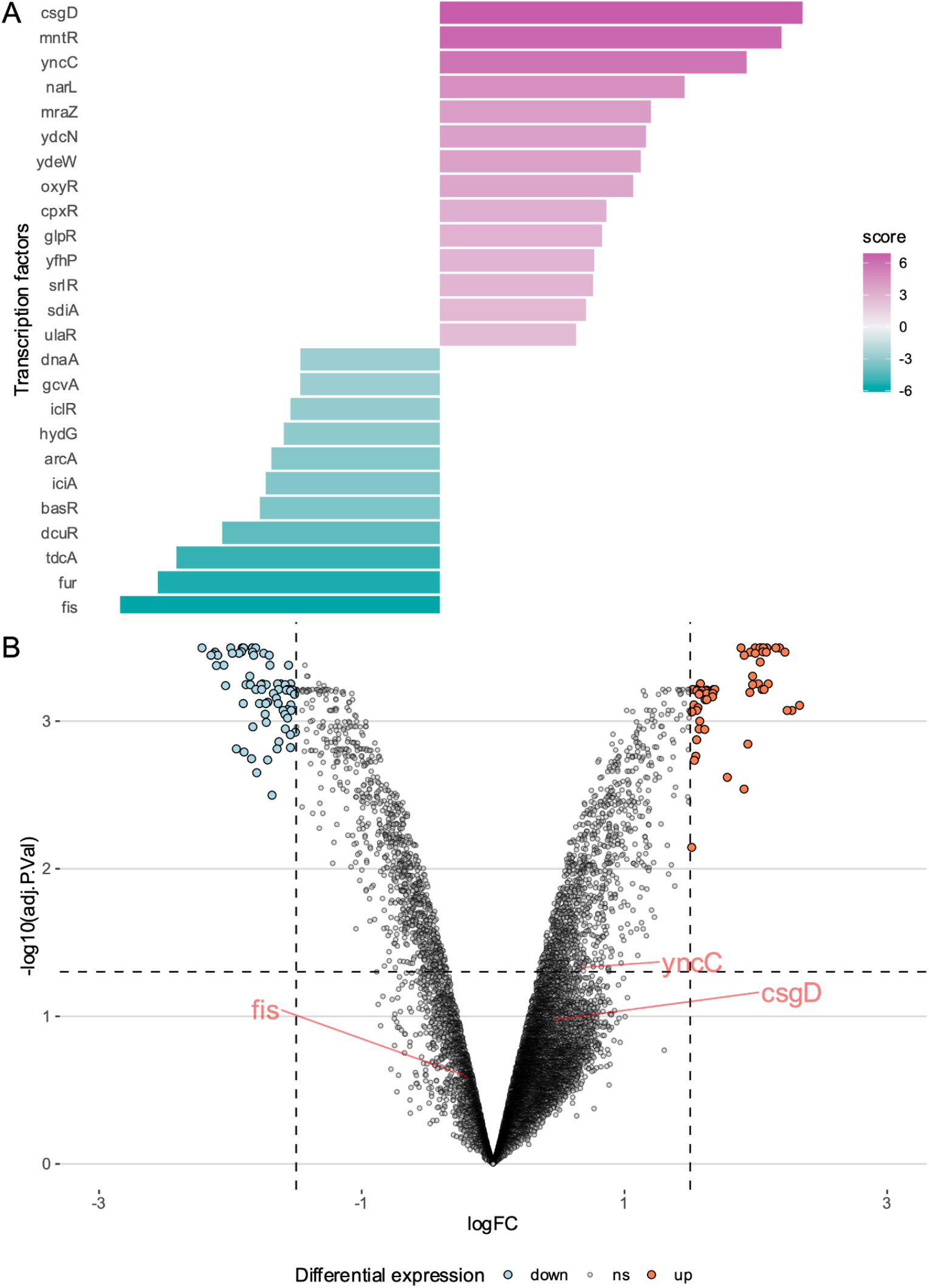
Transcription factor activity of *Salmonella* entering the stationary phase. A: Top ranked transcription factor activities based on differentially expressed genes between the stationary and log phase cultures. B: The most active and inactive transcription factors; CsgD, YncC and fis are labelled. Although these TFs are not differentially expressed themselves, the collective behaviour of their target genes suggest activation (or inactivation).

## Discussion

In this work, we have created a novel resource to infer transcription factor activity in *Salmonella*. The goal of this extension to SalmoNet2 is to bring more functional analysis tools available for the study of model organisms to the *Salmonella* community. Combining transcription factor activity inference with other recently introduced functional analysis tools to the *Salmonella* field, such as pathway enrichment (*25*), has the potential to assist in the characterisation and a greater understanding of transcriptomics experiments. Utilising transcription factor activities could aid antibiotic exposure profiling by inferring TF activity shifts following antibiotic stresses; better understanding of pathogen responses to host environmental stresses such as oxidative or bile stress; while dual-RNA-seq studies could highlight how *Salmonella* TF activities correlate with host pathways.

By determining the activity levels of transcription factors from transcriptomics data, we have a chance to interpret the system in a way that both reduces the dimensionality of transcriptomics data, and increases its robustness, as the activity measure of the individual transcription factors gets determined by the collective behaviour of their target genes (*26*) rather than single data points. To help interoperability with existing tools, we created SalmonAct in the format specifically required for decoupleR, a tool for activity estimation from ‘omics data. Naturally, the resource can also be utilised with other applicable standalone activity inference approaches, such as VIPER (*27*). We also provide helper scripts and tutorials on the SalmoNet2 website to help interested parties analyse their data with both sets of tools.

The main limitation of our approach is that the predictive power of the method is determined by the quality and coverage of the underlying PKN. Our novel PKN consists of interactions from available regulatory databases, interolog transfer from *E. coli*, and an extensive source of regulatory perturbation experiments from SalComRegulon. Regulogs may be appropriate for *Salmonella* strains with the greatest synteny to *E. coli*, but may also be a limiting factor in the case of strains that have undergone extensive pseudogenisation in many of their genes, as is the case for many extraintestinal *Salmonella* strains as they adapted to an invasive lifestyle (*3*). The predictive power of SalmonAct can nonetheless be improved in the future as more relevant datasets become available that elucidate the behaviour and targets of individual transcription factors in *Salmonella*.

## Methods

### Establishing the SalmonAct PKN

To establish the PKN we combined signed, directed regulatory interactions from a number of sources:

1. PRODORIC and CollecTF: Experimentally validated, *Salmonella*-specific signed and directed transcription factor - target gene interactions were included from the PRODORIC and CollecTF databases (*12, 13*).
2. *SalComRegulon (15)*: *Salmonella*-specific interactions were inferred from the SalcomRegulon dataset, a transcriptomics compendium containing 18 transcription factor knockouts of *S*. Typhimurium 4/74. The perturbation data deposited here was used to determine the inhibitory or stimulatory nature of the transcription factor knockouts, and their regulated target genes. A |log2FC| >= 3 cutoff of the responding genes was used as a threshold for inclusion. As the SalcomRegulon data contains regulatory knockouts, we determined their mode of regulation based on the direction of change following the knockout, i.e. log2FC >= 3 were encoded as inhibitory (-1), while log2FC <= -3 were considered activatory (+1).
3. RegulonDB: Information was retrieved from RegulonDB (*28*), a database of transcriptional regulation in *Escherichia coli* K-12. These conserved interactions were imported based on the concept of regulogs (*29*), using the protein orthology relationships established for SalmoNet2 (*16*).
4. RegPrecise: manually curated, signed TF - target gene relationships were downloaded from the RegPrecise website (*14*). Interactions with no annotated mode of regulation were removed.

We then harmonised the confidence levels from the resources (Confirmed, Strong, Weak) ensuring consistent annotation across the network, to help users fine-tune their search terms. Confirmed *Salmonella*-specific interactions had robust experimental validation from CollecTF or PRODORIC. RegulonDB regulog interactions were allocated into Strong and Weak categories, based on the quality of *E. coli* interactions, as established by RegulonDB initially. Most RegPrecise interactions were noted as “Strong”, however, for TFs with an ambiguous mode of regulation the primary annotated mode was used, and these associations were categorised as “Weak” due to this ambivalence. Interactions mapped from SalComRegulon perturbation data were assigned into the “Weak” category, as the perturbations do not capture the differences between primary or secondary order regulatory effects caused by the transcription factor hierarchy present in the cell; these edges are sign-consistent hypotheses, not direct binding evidence. Interactions present in multiple resources were collapsed, and the sources listed in the corresponding column. In case of partially matching interaction data, the higher quality interactions were kept. In cases where the partially matching interactions were in the same category (Weak) the more *Salmonella* specific data sources were kept (RegPrecise & SalComRegulon > RegulonDB). From the initially assembled PKN, orthology-based mapping was done for all strains included in SalmoNet2, using the orthologous relationships established with OMA (*30*) for SalmoNet2. We provide a mapping table on the project github page to enable availability for all 20 strains in the database. The target genes in the mapped resources were noted using strain specific locus tags.

### Inferring transcription factor activity of *Salmonella* regulons

Differential expression for the selected transcriptomics data sets (GSE35938, GSE11486) was established using GEO2R with default settings. In the GSE35938 dataset wild type strains grown in LB were contrasted to *ompR* mutants, while in the GSE11486 dataset wild type strains grown in log phase were compared to wild type strains grown in stationary phase.

The R package decoupleR (version 2.10) was used to infer the activities from the PKN (“Strong” and “Confirmed” interactions) and transcriptomics datasets. The univariate linear model (ULM) approach was used to determine transcription factor activity, with the minimum size of target sets set to 5, and a significance cutoff of 0.05.

The ULM approach creates a linear model where the predictor variable is the interaction weight between the transcription factor (TF) and each gene (-1 or +1 based on the mode of regulation included from the sources the network was built from), and the response variable is the expression level of the genes. This way, ULM evaluates how well the interaction weights of a TF predict the expression of its target genes, by fitting a linear regression resulting in a slope value that represents the relationship between the TF’s interaction weights and gene expression. A positive slope means that as the interaction weight increases, the gene expression also tends to increase, indicating the TF is active. Conversely, a negative slope would suggest that the TF is inactive. The score for each TF’s activity is derived from the t-value of the slope, which indicates the strength and significance of the relationship. A high positive t-value means strong activation of the TF, while a high negative t-value indicates inactivity. While the ULM method was used in these examples, decoupleR can perform other approaches aimed at TF activity estimation with the same input data, such as VIPER or AUCell (*27, 31*), and can provide a consensus score as well utilising all 11 methods (returning a mean z-score across methods).

P-values were adjusted using the ‘fdr’ method using the p.adjust function in R. R scripts to replicate the differential expression results and transcription factor activities can be found in the project github repository: https://github.com/korcsmarosgroup/salmonella-TF-activity.

## Data availability

The SalmonAct PKN is available on the project github repository: https://github.com/korcsmarosgroup/salmonella-TF-activity and the SalmoNet2 website (http://salmonet.org).

## Acknowledgements

T.K. was supported by the NIHR Imperial Biomedical Research Centre Organoid Facility, and the UKRI BBSRC Gut Microbes and Health Institute Strategic Program BB/R012490/1 and its constituent projects BBS/E/F/000PR10353 and BBS/E/F/000PR10355, as well as a UKRI BBSRC Core Strategic Program Grant for Genomes to Food Security (BB/CSP1720/1) and its constituent work packages, BBS/E/T/000PR9819 and BBS/E/T/000PR9817. B.B and T.K. were also supported by the UKRI BBSRC Institute Strategic Programme Food Microbiome and Health BB/X011054/1 and its constituent project BBS/E/F/000PR13631. RK was supported by BBSRC Institute Strategic Programme Microbes and Food Safety BB/X011011/1 and its constituent projects, BBS/E/F/000PR13635 and BBS/E/F/000PR13636.

## Supplementary Materials

**Table S1:**
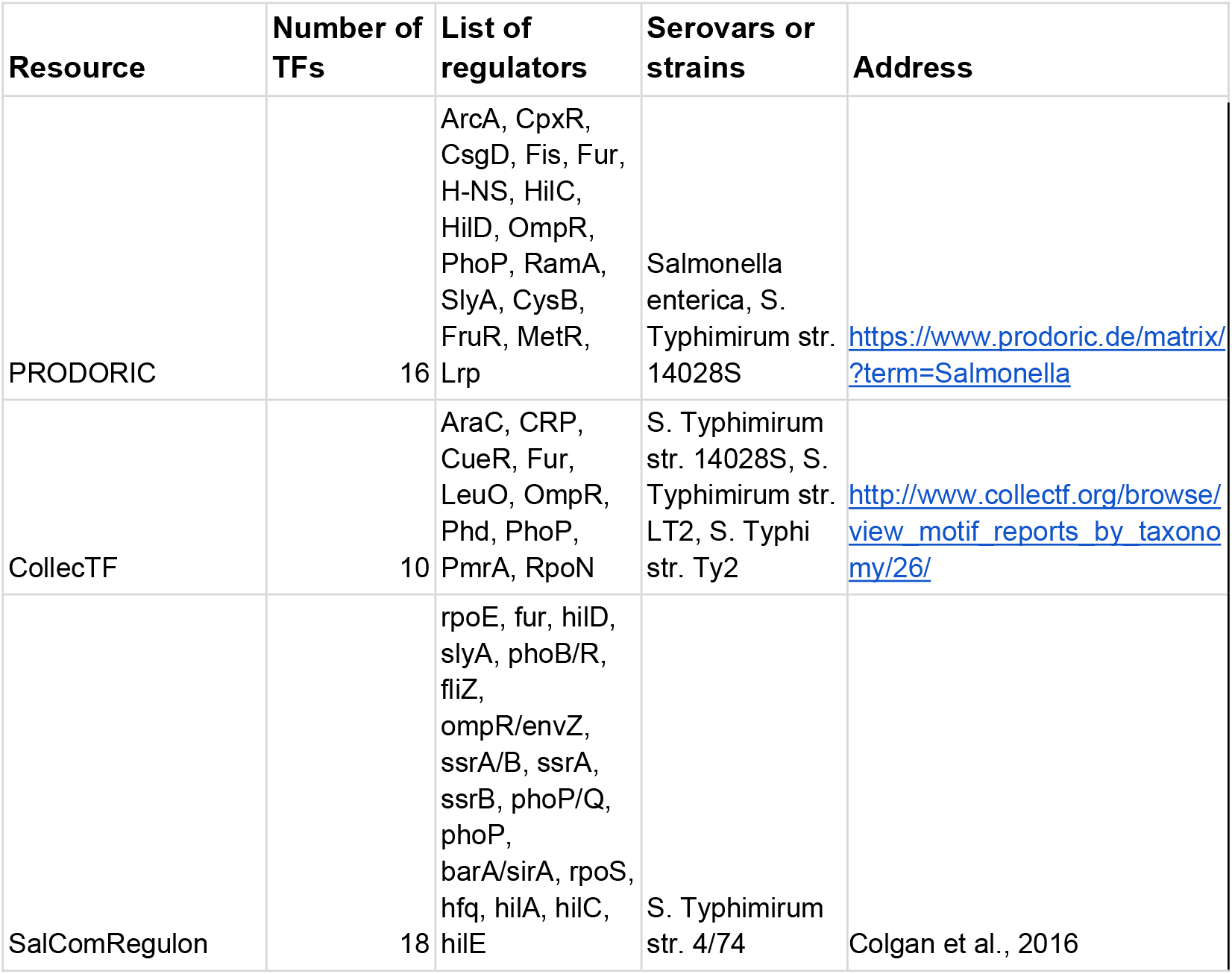
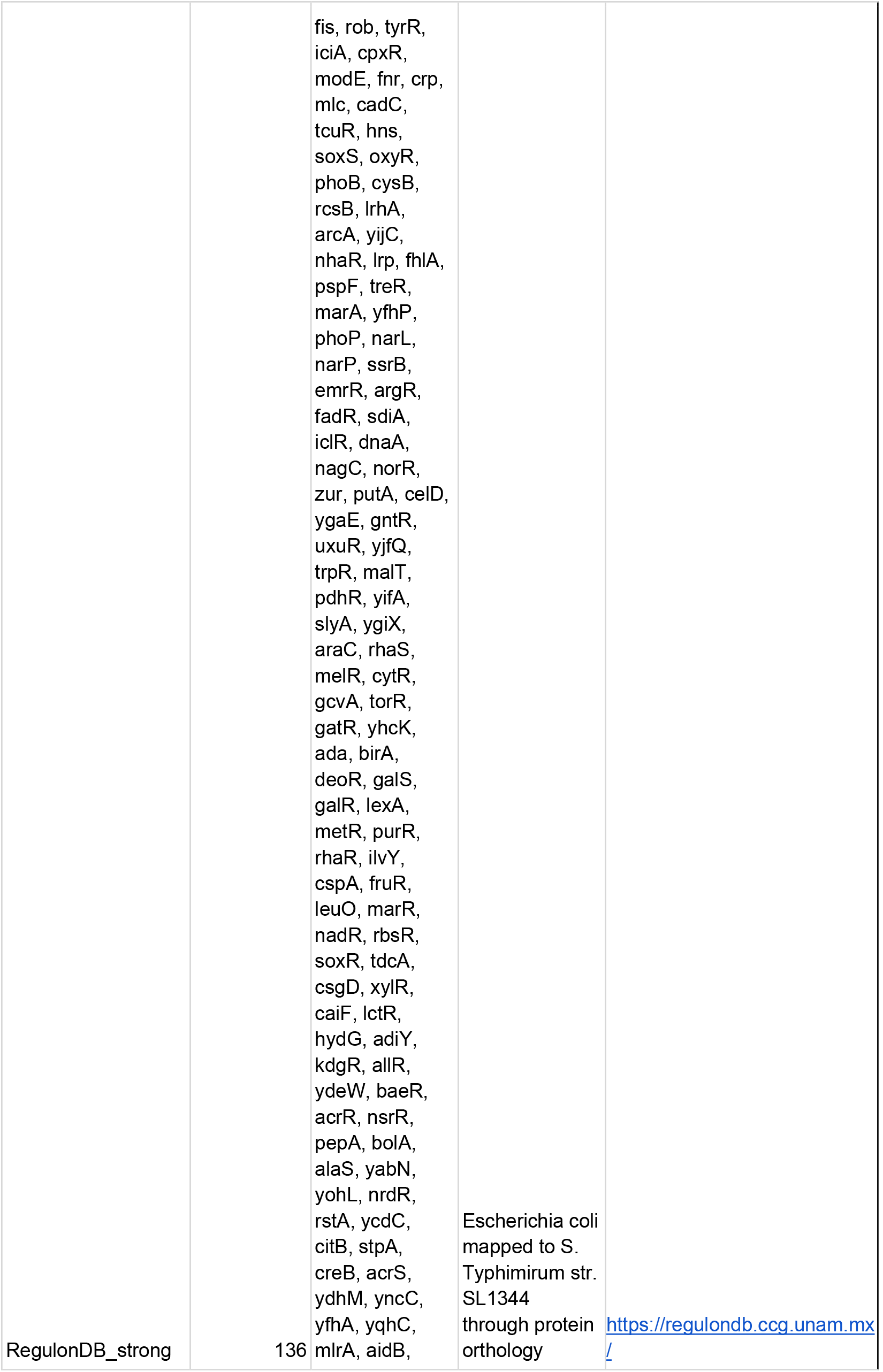
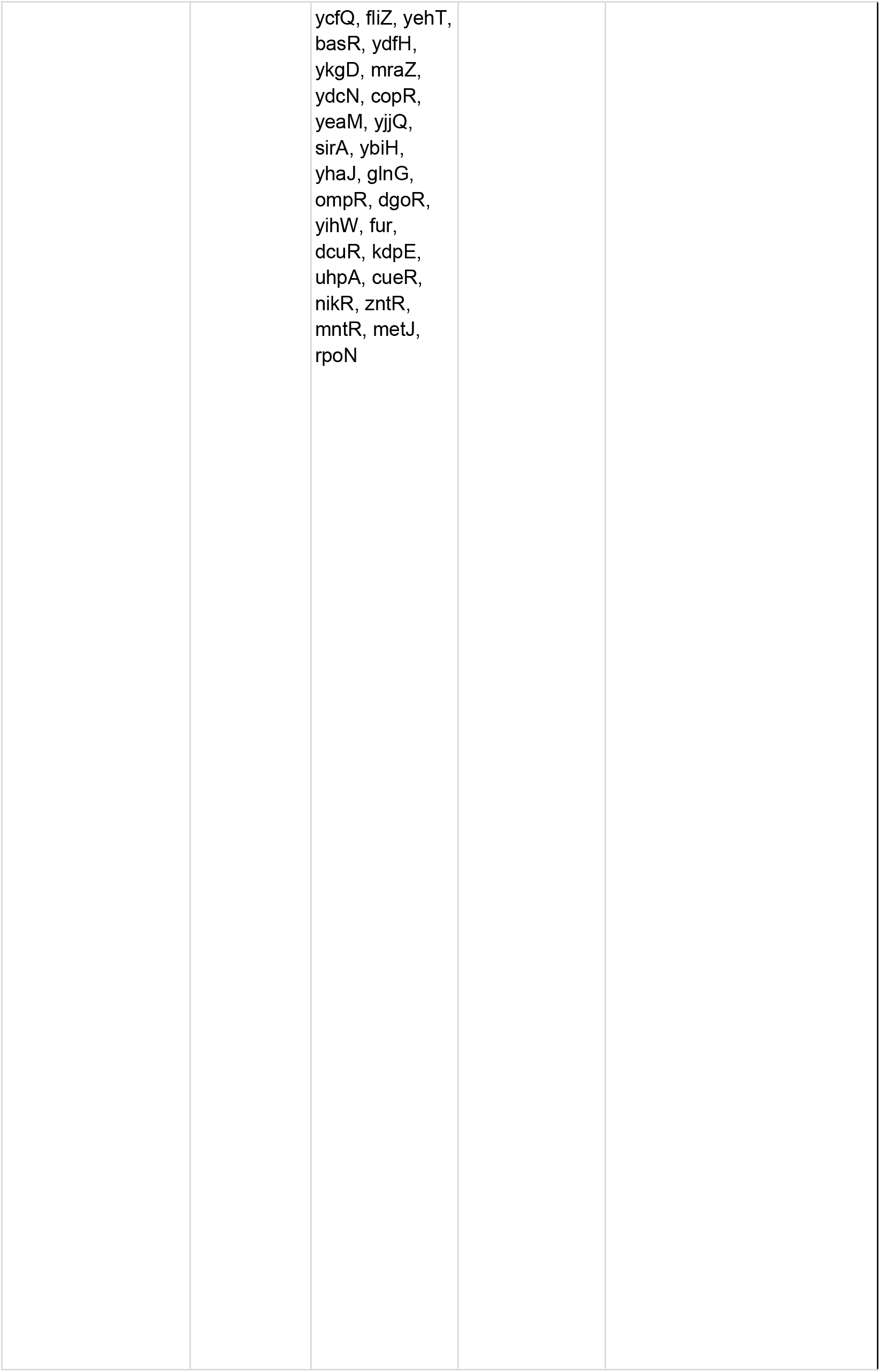
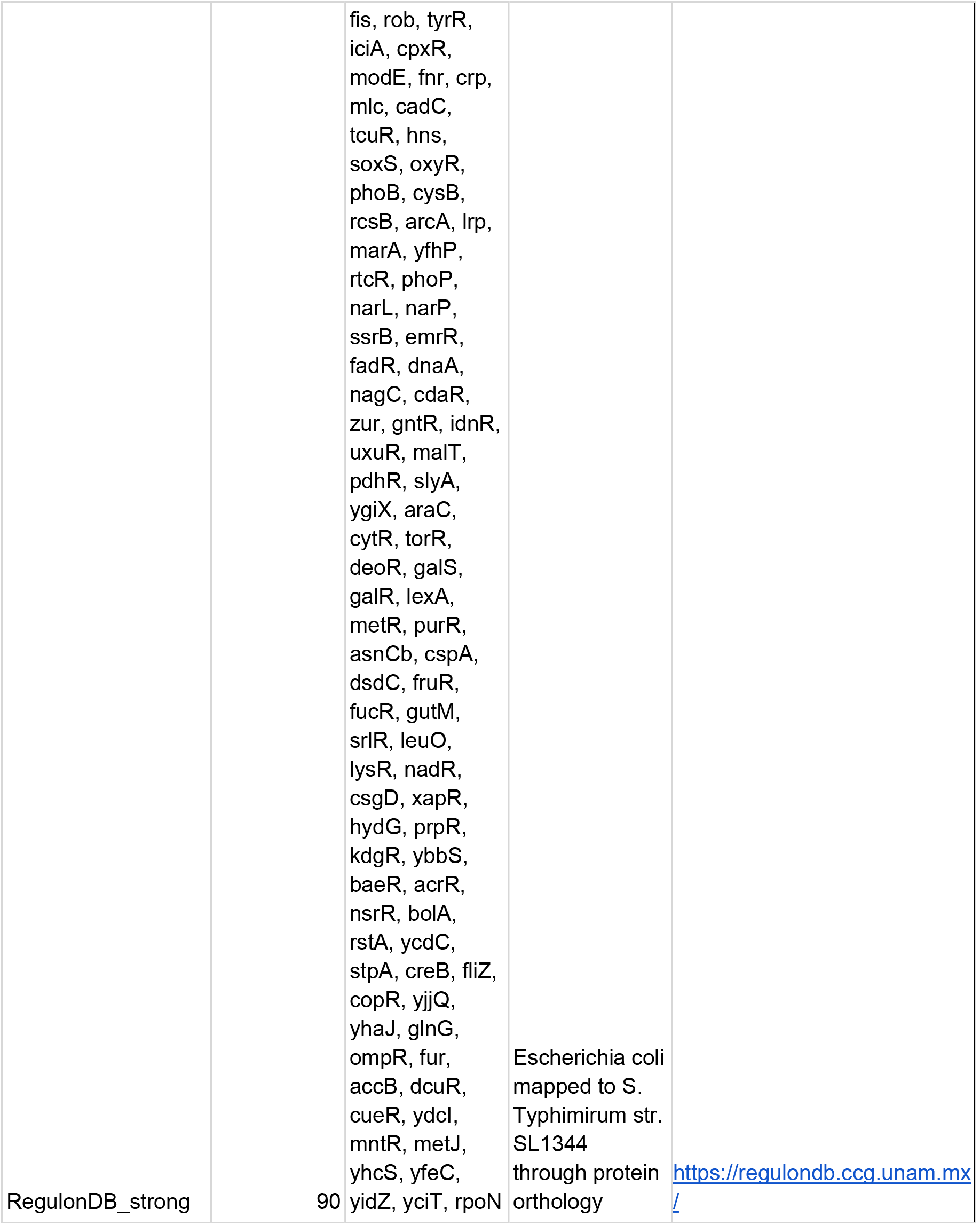
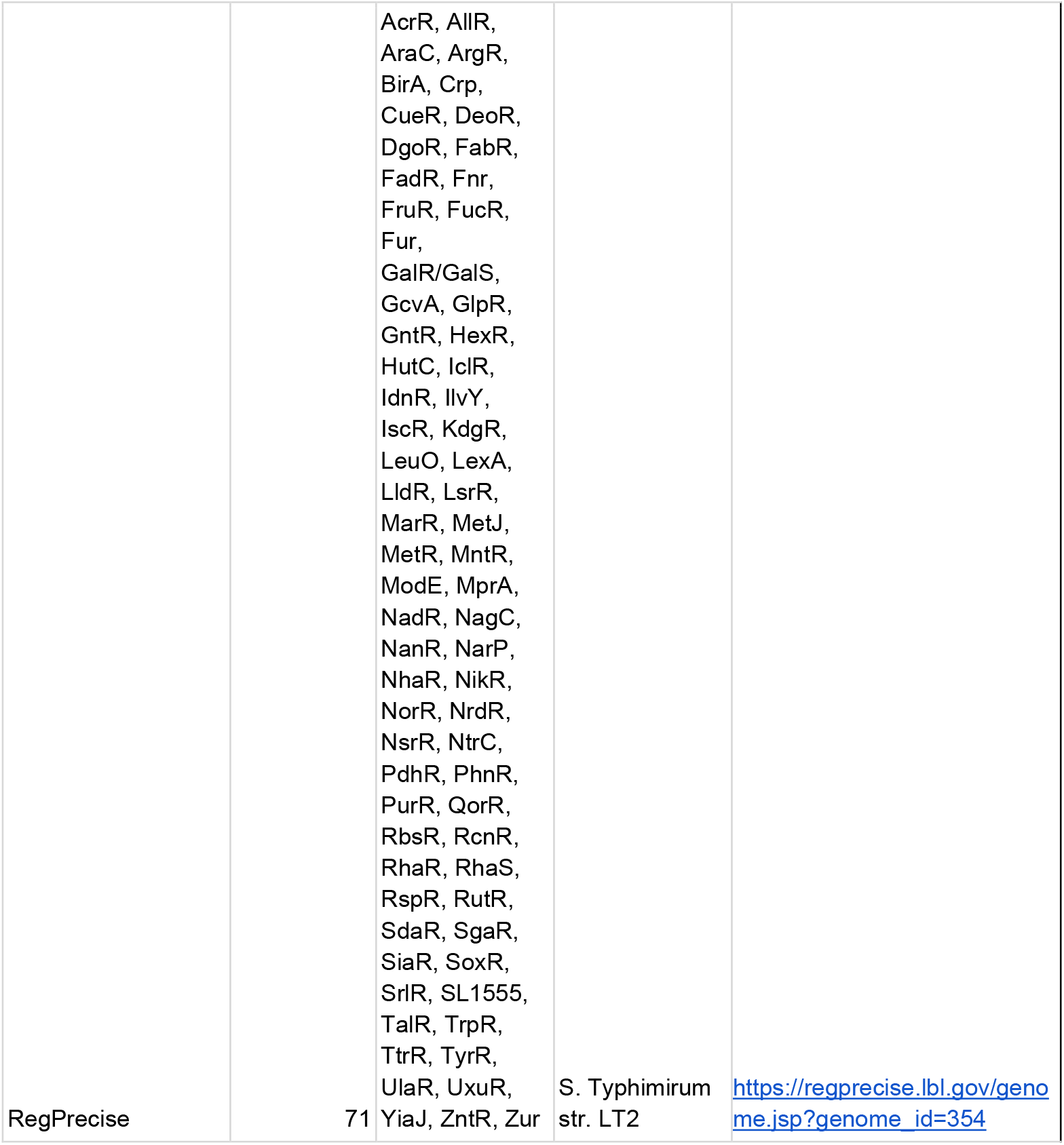
List of resources, transcription factors and source serovars or strains used to construct the PKN.

